# The interplay of top-down focal attention and the cortical tracking of speech

**DOI:** 10.1101/813147

**Authors:** D Lesenfants, T Francart

## Abstract

Many active neuroimaging paradigms rely on the assumption that the participant sustains attention to a task. However, in practice, there will be momentary distractions, potentially influencing the results. We investigated the effect of focal attention, objectively quantified using a measure of brain signal entropy, on cortical tracking of the speech envelope. The latter is a measure of neural processing of naturalistic speech. We let participants listen to 44 minutes of natural speech, while their electroencephalogram was recorded, and quantified both entropy and cortical envelope tracking. Focal attention affected the later brain responses to speech, between 100 and 300 ms latency. By only taking into account periods with higher attention, the measured cortical speech tracking improved by 47%. This illustrates the impact of the participant’s active engagement in the modeling of the brain-speech response and the importance of accounting for it. Our results suggests a cortico-cortical loop that initiates during the early-stages of the auditory processing, then propagates through the parieto-occipital and frontal areas, and finally impacts the later-latency auditory processes in a top-down fashion. The proposed framework could be transposed to other active electrophysiological paradigms (visual, somatosensory, etc) and help to control the impact of participants’ engagement on the results.

## Introduction

Modelling neural processing of auditory stimuli is an important theme in audiology and neuroscience. While many studies rely on the use of simple, repetitive and non-realistic time-locked stimuli (e.g., clicks), there has been increased focus on the use of naturalistic speech (e.g., see ^1^) as the ecological validity is greater ^2^: the brain may process natural speech differently from artificial repeated stimuli. The interaction between a feature (e.g., the envelope or the phonetic representation) of the presented speech and the recorded brain signals can be modelled using approaches like the temporal response function (TRF ^3^) based on linear regression. This linear transformation can then be used to either reconstruct the speech feature from the brain signals (using a so-called backward model) or predict the brain response across the scalp to a specific speech segment (using a so-called forward model). The correlation between the reconstructed/predicted artificial and the actual time-course is then a measure of cortical speech tracking. Measures of cortical speech tracking quantify speech processing within the brain, opening doors to, for example, an objective measure of individual speech understanding ^4–7^ or auditory attention decoding in a cocktail party scenario ^8–10^.

For the tracking of different speech features, the cortical activity in the delta (1 – 4 Hz) and theta (4 – 8 Hz) frequency bands has been found to be most important ^4,5^. This can be partially explained by the correspondence with the syllable (4 – 8 Hz), word and phrase (1 – 4 Hz) rates of speech, suggesting a real-time transformation of the continuous acoustic in those different linguistic representations ^11–15^. On the contrary, the alpha and higher frequency bands show weak and noisy responses, presumably reflecting the weaker phase locking of EEG neural data to the speech envelope at higher frequencies ^6^.

While the alpha and beta EEG oscillations hardly seem to track speech features, they play a key role in the modulation of attention ^16,17^. Although the term “attention” is widely used in several fields, delimiting and defining this broad term was a tricky problem ^18,19^. Attention is defined by the process of volitionally concentrating on specific information while disregarding other information ^20^. There are four main types of attention ^21,22^: selective attention (i.e., the ability to focus on one out of several stimuli), divided attention (i.e., the ability to maintain focus on more than one stimulus at a time), sustained attention (i.e., the ability to actively focus on a specific stimulus over an elongated period of time), and alternating attention (i.e., the ability to switch back and forth between tasks).

Several studies have investigated the effect of attention on auditory brainstem responses (ABRs), auditory steady-state responses (ASSR ^23^), or late auditory evoked potentials, mainly using an auditory attention protocol relying on the participant’s selective attention. It remains debated whether attention impacts the ABR, particularly regarding the inconclusive results on the effects of attention on the ABR to short clicks or pure tones ^24–28^. However, this could be explained by the tiny amplitude of the ABR requiring a large number of repetitions of an identical, short and meaningless stimulus leading to a decrease in attention. Recently, Forte et al. (2017) measured the ABR to running speech and showed that the ABR can be modulated by attention ^29^, most likely in a top-down fashion. At the cortical level, Linden et al. (1987) did not find an effect of selective attention on the 40 Hz-ASSR ^30^. The impact of attention on lower ASSR frequencies is controversial: Müller et al. (2009) found some differences for the 20 Hz-ASSR ^31^ while Skosnik et al. (2007) did not ^32^. Studies using late auditory evoked potentials have shown that responses in cortical regions are modulated by attention ^33–37^. In particular, many researchers investigated the effect of selective attention in auditory attention decoding studies and showed attentional modulation of speech tracking. However, the latency of the attentional modulation effect has been controversial: Ding and Simon (2012a, 2012b) showed an early attentional gain effect at a time lag around 100 ms ^11,38^, while others reported a longer-latency attentional effect above 150-ms time lag ^14,39,40^. Following this research, we could hypothesize that only later responses with a cortical origin are modulated by attention. However, these studies focused solely on <u>selective</u> attention and little is known about the impact of changes over time in the level of <u>sustained</u> attention in the modelling of cortical speech tracking.

Attention is a conscious, active process and has a robust top-down effect on processes along most of the cortical auditory pathway ^29,34^. A network of frontal and parietal cortical areas is involved in the selection required for top-down attention ^41–45^. In the following, the level of sustained attention during a task will be referred to as “focal attention” or “attention”. Understanding and accounting for the user’s attention is important in active paradigms and could help understanding the fluctuation of cortical tracking over time. However, it is difficult to behaviorally measure the level of focal attention for each participant since it would extend experiment time and require an efficient behavioral scale. Objective measures of focal attention using evoked brain responses have been proposed, such as the brain response at a specific frequency ^46^, or energy changes of beta ^47^ or alpha ^17^ EEG rhythms. Lesenfants et al. (2018) showed that focal attention could be tracked using spectral entropy measures ^48,49^. This measure could help distinguish active from passive periods during an active task in both healthy volunteers and participants with locked-in syndrome (i.e., fully conscious but suffering from brain injuries leading to full or partial tetraplegia).

We here aim to study the impact of <u>sustained</u> attention, objectively evaluated using brain entropy measures, on the modelling of cortical speech tracking and evaluate the potential of correcting for attention in a standard speech paradigm.

## Methods

### Participants

In this study, we included 19 normal hearing volunteers (one male; age 22 ± 4 years; two were left-handed). All the participants were Flemish-speakers and their hearing was evaluated by pure tone audiometry (thresholds lower than 20 dB HL for 125 Hz to 8000 Hz using a MADSEN Orbiter 922–2 audiometer). The experiment was approved by the Medical Ethics Committee UZ KU Leuven/Research (KU Leuven, Belgium) with reference S59040 and all participants provided informed consent. All methods were carried out in accordance with relevant guidelines and regulations.

### Experiment

Each participant sat comfortably and wore Etymotic ER-3A insert phones (Etymotic Research, Inc., IL, USA), protected from electromagnetic fields with CFL2 boxes from Perancea Ltd. (London, UK). The room, in which the experiment took place, was electromagnetically shielded and soundproofed. Prior to the experiment, we calibrated the acoustic system using a 2-cm^3^ coupler of the artificial ear (Brüel & Kjær, type 4192, Nærum, Denmark). They were instructed to listen attentively to the children’s story “The Little Mermaid”, written and narrated in Flemish by Katrien Devos. Every 15 minutes, there was a short break and a question was asked about the contents of the story, to ensure compliance. The stimulus was 44 minutes long and was presented binaurally at 60 dBA using the APEX 3 (version 3.1) software platform (ExpORL, Dept. Neurosciences, KU Leuven, Belgium; see ^50^) and an RME Multiface II sound card (RME, Haimhausen, Germany). During the presentation of the speech stimulus, the experimenter sat outside the room and visually inspected the EEG signals in real-time.

### Recordings

64-channel EEG was recorded at a sampling rate of 8192 Hz using Ag/AgCl ring electrodes and a Biosemi ActiveTwo system (Amsterdam, Netherlands). The electrodes fixed to the cap were positioned according to the International 10-20 system.

### Data analysis

All analyses were done with custom-made Matlab scripts and the mTRF Toolbox ^3,51^.

#### Data preprocessing

EEG artifacts were removed using a multi-channel Wiener filter algorithm ^52^. We then re-referenced each EEG signal to a common average reference before downsampling from 8192 to 1024 Hz to decrease processing time. The speech envelope was extracted using a gammatone filterbank (24 channels spaced by 1 equivalent rectangular bandwidth, with center-frequencies from 50 Hz until 5000 Hz) followed by a power law ^53^, and then downsampled to 1024 Hz.

#### Attentional level

Entropy measures ^48^ were extracted from pre-processed EEG signals using power spectral estimation based on Multitapers Spectral Analysis (7 tapers) and computed using the following formula:

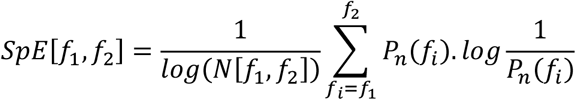

where:

- P_n_(f_i_) is the normalized power spectrum at frequency f_i_. The normalization process divides each frequency power by the sum of all frequency powers.
- N[f_1_,f_2_] is equal to the total number of frequency components in the range f_1_-f_2_.

This measure reflects signal regularity or predictability; usually, a high value corresponds to low regularity or predictability (e.g., uniform distribution of the power spectrum), while a low value corresponds to high regularity (e.g., concentration of the activity in a single band). Importantly, while this measure is classically computed in the frequency band from f_1_ = 0.5 to f_2_ = 32 Hz, we here decided to reduce this band to 8.5-32 Hz, with a 0.5 Hz step. This limits the influence of the speech processing, investigated in the delta and theta EEG bands, on this measure of attention. Since attention regroups a fronto-parieto-occipital network ^41–45^, we can hypothesize that mainly electrodes over the frontal (F), parieto-occipital (PO) and occipital (O) areas will be involved in this mapping. This was confirmed by our prelude analysis, detailed in the Appendix. In the following, the averaged spectral entropy over the frontal and the parieto-occipital electrodes will be used as a marker of the level of attention over time.

#### Cortical speech tracking

Pre-processed EEG signals and speech envelope were bandpass filtered between 0.5-8 Hz using zero phase Butterworth filters with 80 dB attenuation at 10 % outside the passband. Stimulus representations and EEG signals were then downsampled to 256 Hz. Each 44 minutes dataset was split into consecutive non-overlapping 1-minute periods and the averaged entropy values were computed for each segment in the frontal and parieto-occipital areas. The data was split in contiguous training and testing sets in a cross-validation paradigm (see Figure 1). Forty minutes were used for training and the remaining 4 minutes for testing. Both the training and the testing sets were then divided in 1-min segments, which were assigned to two classes: the first class (“high SpE”) contains the 50% highest entropy epochs while the second (“low SpE”) contains the 50% lowest entropy epochs. This resulted in a “high SpE training” (20 1-minute epochs), “low SpE training” (20 1-minute epochs), “high SpE testing” (two 1-minute epochs) and “low SpE testing” (two 1-minute epochs) dataset. For each subject, we first normalized (parametric normalization) both the speech envelope and EEG signals of the training set, prior to finding a mapping between the speech envelope and the recorded EEG data, commonly referred to as a decoder. A decoder aims to optimally reconstruct the speech envelope by linearly combining EEG electrode signals and their time shifted versions (a stimulus evokes neural responses at different delays along the auditory pathway). During the training step, each time-delayed channel’s weight is determined. The modelling of the brain-speech interaction can then be seen as the computation of a mapping g(n,τ), where *“n”* is an EEG channel and “τ” is a specific time delay between the speech and the neural response, so:

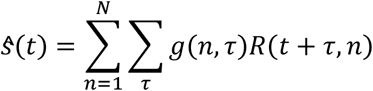

**Figure 1.**
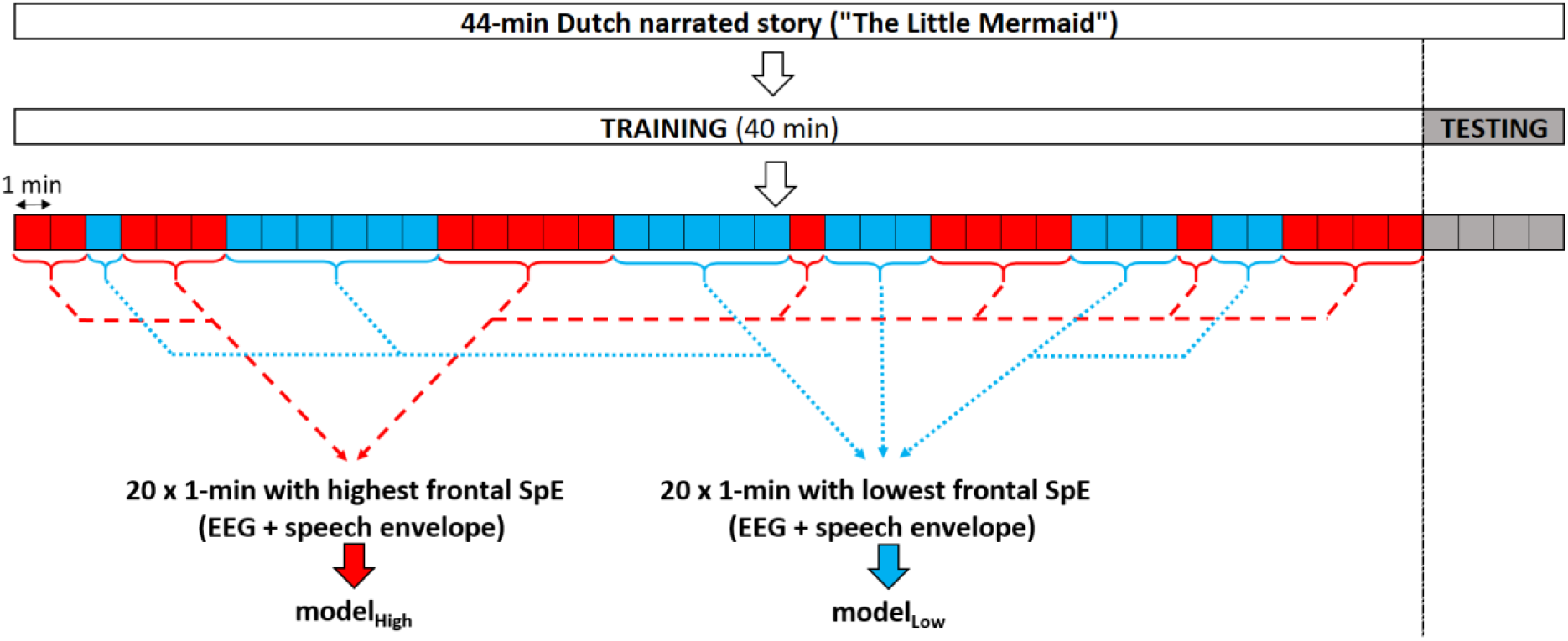
Segmentation of the dataset. In red/blue, example 1-minute epochs with a high/low averaged attentional level, computed using spectral entropy values in the frontal and parieto-occipital areas are shown.

in which “g” can determined by minimizing a least-squares objective function:

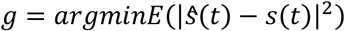

In the previous equations, *ŝ*(*t*) and *s*(*t*) are the reconstructed and the actual speech envelope respectively; *E* denotes the expected value; R is a matrix containing the time-shifted neural responses of each channel. In practice, the linear decoder is computed by solving *g* = (*RR^T^*)^−1^(*RS^T^*), which can be done using ridge regression on the inverse autocorrelation matrix. In this study, a subject-specific backward model between the speech envelope and the EEG was computed using ridge regression with a time-lag window of 0-400 ms using either the low SpE training or the high SpE training dataset. This model was used to reconstruct the speech envelope using the EEG signals from the test set. The cortical speech tracking was computed as the correlation (Spearman’s rank-order correlation) between the actual and reconstructed speech envelope.

The estimation of a temporal response function is similar to the calculation of the decoder, but we now predict EEG instead of reconstructing the envelope (i.e., it goes the other way around and works per channel). In this study, we decided to work with backward models (i.e., “decoders”) to investigate how stimulus features are encoded in neural response measures because no pre-selection of electrodes is required ^54^ and this method is more sensitive to interesting signal differences in highly correlated channels because of its multivariate properties ^3^. However, backward model parameters are not readily neurophysiologically interpretable ^55^. Therefore, we separately calculated the forward model to investigate model parameters across the scalp. The temporal response functions shown in Figure 5 are modelHigh and modelLow of Figure 1 but in the forward domain and averaged across folds and subjects (by averaging the covariance matrices of the EEG signals ^56^).

#### Statistics

A permutation test ^57,58^ was used to evaluate the significance level for each model (1000 repetitions, p < 0.01). The significance of change between conditions was assessed with a non-parametric Wilcoxon signed-rank test (WSRT; α = 0.01), with Bonferroni correction for multiple comparisons. Finally, we investigated whether the temporal response functions of the “high” and “low” attention condition were significantly different using a cluster-based analysis (α = 0.01) ^59^. This approach solves the multiple comparison problem by clustering adjacent lags and channels, and allows indicating which latencies and channels contributed most to this difference.

## Results

When considering stimulus reconstruction accuracy, both the data used for training the decoder and for testing the decoder can play a role. We first investigated the effect of training data. Using only periods of higher attention to train the decoder, the median cortical tracking across subjects reached a correlation of 0.22 ± 0.07 (median ± iqr) at group-level (see Figure 2, left panel, in red). Using only periods of lower attention to train the decoder, the median cortical tracking was 0.15 ± 0.08. At the individual level, 14/19 individuals presented a higher cortical tracking using periods with high attentional level (WSRT, for each of these 14 individuals p<0.05) than periods with low attentional level. Two out of the 19 subjects showed no difference. Interestingly, comparing between training the decoder on the whole training set versus only the high-attention segments, 12/19 participants achieved higher cortical tracking using only periods of higher attention (WSRT, for each of these 12 individuals p<0.05), even though there is only half as much data in the high-attention part. Two out of the 19 subjects showed no difference. This suggests we could decrease the session time by tracking the participant’s attention during the recording (e.g., by comparing in real-time the level of attention in every new minute of recording, while the first minute is set as a baseline under the assumption that the participant is attentive on the task straight at the beginning of the experiment).

**Figure 2.**
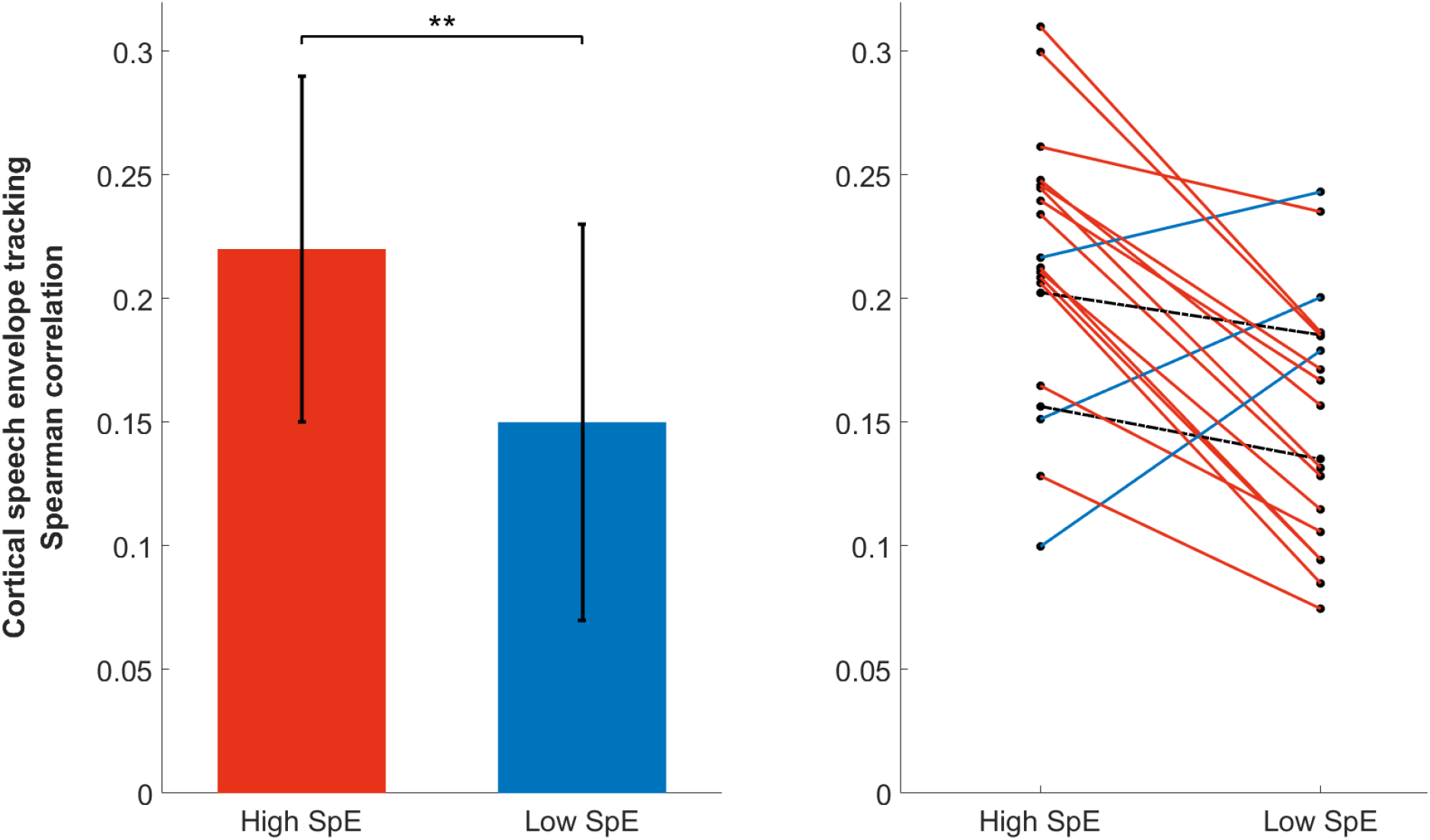
Group (left) and individual (right) Spearman correlation using only periods of high (high SpE) or low (low SpE) attention on the computation of the decoder (median ± iqr). Note that more than 80% of the group showed higher correlation using a decoder based on periods of high attention than low attention. Each dot represents the cortical speech tracking for a participant (i.e., the cortical speech tracking averaged over folds). For each fold, the cortical speech tracking is computed over a complete testing period (i.e., 4-minutes of data). This means that each Spearman correlation value is computed by comparing 61440 data points (i.e., 4×60×256) of the actual and reconstructed speech envelope. All the correlations are significant (a permutation test was used to evaluate the significance level for each model, which was at around 0.03 for each participant).

Next, we investigated the effect of testing data. Using the same decoder, we did not observe any difference in the cortical tracking computed using the reconstructed envelope in either testing periods of higher or lower attention (see Figure 3; the left and right panel show individual performance using a decoder trained on periods with, respectively, higher and lower attention only).

**Figure 3.**
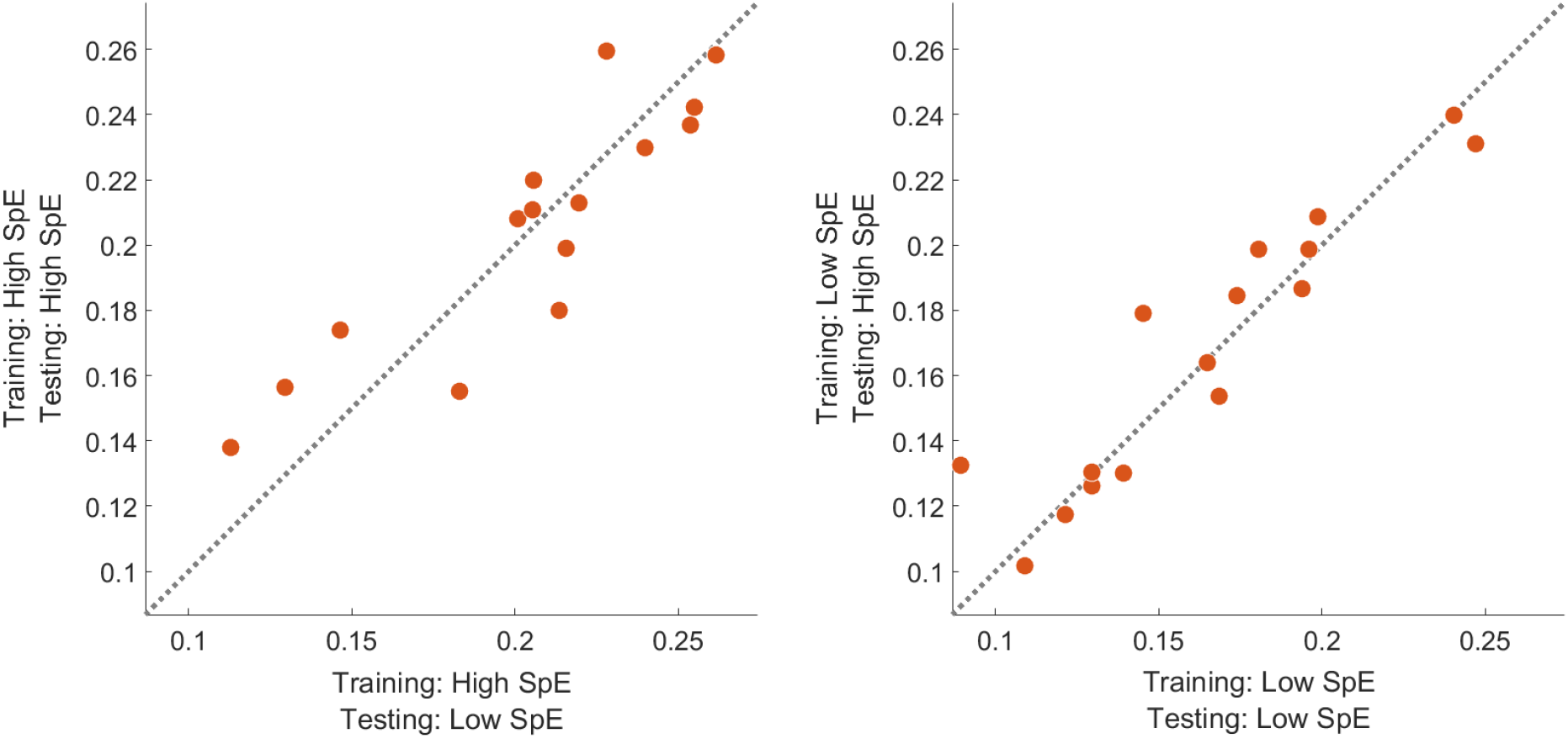
Individual performance using either a training set based on periods of high (left panel) or low (right panel) entropy and a testing set based on periods of high (y-axis) or low (x-axis). Note that while an improvement could be observed when the decoder is trained using periods of high entropy as compared to low entropy, the selection of periods in the testing set did not affect the performance.

We then compared forward models built using periods of either low or high level of attention. For the early latencies (below 100 ms), no delay could be visually observed between the temporal response function built using either high or low attention periods. For latencies between 100 and 300 ms, a delay could be visually observed in the left and right frontal and fronto-central channel locations (see Figure 4, electrodes F and FC). This delay is around 8 ms (mainly at FC1, FC3 and FC5 channels) at 100-ms latencies and increased to 16 ms for latencies between 200 and 300 ms. For later latencies (i.e., above 300 ms), no delay could be observed. A cluster-based analysis allows showing a significant difference in TRFs between the high and low attention condition. Figure 4 (panel B) shows latencies and channels contributed most to this difference. This difference is most pronounced from 242 to 290 ms. For visualization purposes, we averaged the TRFs across channels (see Figure 4, panel C). The difference results from a 15.7 ms delay (see Figure 4, panel C).

**Figure 4.**
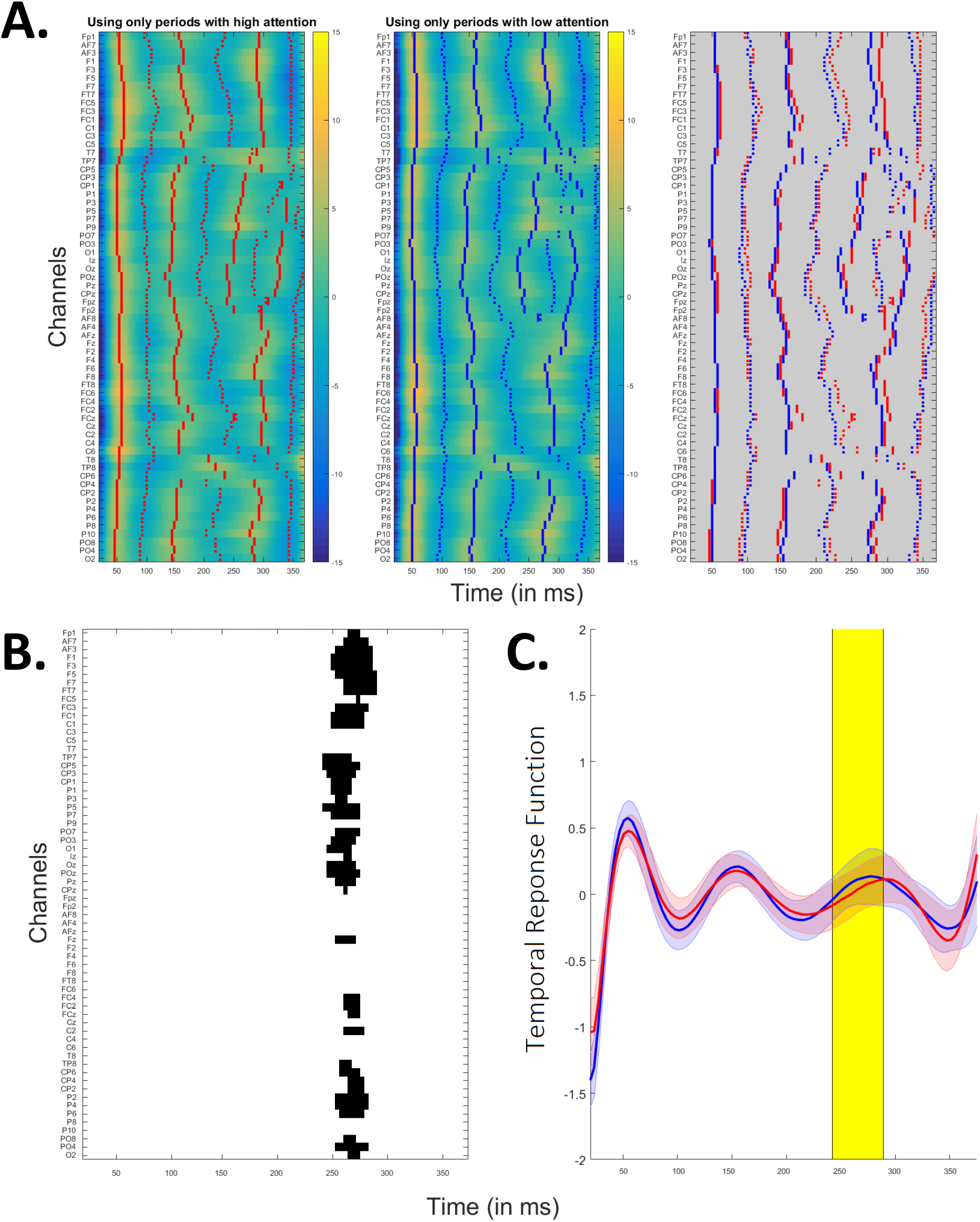
Panel A: Temporal response function using periods of high (left panel) or low (center panel) attention for training the model. In red/blue, the peaks are indicated. In the right panel, peaks from the two different conditions are overlaid. Note the delay in later latencies (between 100 and 350 ms) observed in the frontal and fronto-central electrodes, while earlier and later latencies did not show any delay in the activations. Panel B: Temporal response function’s differences at each latency. Channels and latencies that were found to contribute significantly to this difference in a cluster-based analysis are indicated in black. Panel C: Temporal response functions using periods of high (red) or low (blue) attention for training the model averaged over electrodes. The TRFs are significantly different, and the difference is most pronounced from 242 to 290 ms (in yellow). The difference mainly results from the peak’s delay and not the peak’s amplitude change.

## Discussion

In this study, we showed that we could improve the modelling of cortical speech tracking using only data from periods with high attention, measured using spectral entropy. Three quarters of the participants showed an improved cortical speech tracking using only 20 minutes of data with high attentional level. Interestingly, they also showed improved performance compared to the use of the complete 40 minutes training dataset. This illustrates the impact of shifts or declines of the participant’s attention in active EEG paradigms.

We first evaluated the impact of the spectral entropy on cortical speech tracking and showed that the modulation of attention in the frontal and parieto-occipital areas impacts the cortical tracking of the speech envelope. This is consistent with many studies that evaluated the interaction of brain networks in the processing of attention and showed increased activity with attention in the frontal and parieto-occipital cortex, as well as in the thalamus, cingulate cortex and temporal parietal junction ^60–62^. The entropy measure requires no assumption about the frequencies that dominate the attentional process. Alternatively, attention could be extracted using alpha power, beta power, a ratio of the two, or others. While our analysis may work using only alpha power, Lesenfants et al. (2018) showed that the entropy measure outperforms the alpha-only rhythms in tracking attention during an active task ^49^.

While focusing on periods with higher attention in the training set allows to improve the modelling of the brain-speech interaction, no clear impact of the testing dataset could be observed in our study, i.e., applying the same decoder on periods of higher or lower attention of the testing dataset did not produce significant differences. This could be explained by an insufficient amount of testing data to find significant effects. Note that, while the differences are not significant, there was a trend of improvement of lower correlations (see Figure 3, left panel, the three leftmost points) using only testing periods of higher attention. This suggests that, while the selection of training periods improves the cortical speech tracking overall, the selection of testing periods corrects for lower performance only.

The evaluation of the temporal response functions resulting from the modelling of the brain-speech interaction using periods with either low or high attentional level highlights the importance of later-latency brain responses, appearing between 100 and 350 ms. We found slightly delayed brain responses to speech in this latency range over the frontal and fronto-central areas. This suggests a top-down frontal attention mechanism during task processing, similar to the P3a event-related brain potential ^63^. It is important to note that the attention mechanism results from a top-down process ^34^, thus we expect this to mainly affect brain responses with a cortical origin (i.e., the later auditory evoked potentials). In figure 5, we schematically illustrated how information could be transferred between the different brain areas. The presentation of the speech to the participant’s ear induces cortical tracking in the early auditory areas (see P50 and N100 peaks in Figure 4). As soon as the speech stimulus reaches the first auditory layers, a bottom-up signal is sent to the parieto-occipital and frontal cortices, inducing a top-down response from each of these areas. The parieto-occipital top-down control here preceeds the frontal top-down control (respectively the red and green circles in Fig.1 of the Appendix). Each of these areas are major components of the dorsal and ventral attentional networks ^64^. Thanks to the direct connections for dorsal-ventral interactions, we could hypothesize that the activation of the frontal area is due to both the bottom-up signal initiated in the auditory area and the putative intra-network connections between the frontal area and the parieto-occipital area, activated earlier (i.e., the green circle preceeds the red ones). Altogether, this allows to reinforce the speech tracking in the later auditory areas (see brain response after 100 ms in Figure 4) using top-down connections. The delay between the emission of this signal and the effect on speech processing could potentially be linked to the time needed to close the auditory-associative-auditory loop (i.e., the time required for an action potential to reach, be processed and come back from the primary auditory area to each of the associative areas involved in the attentional network). Interestingly, in a review paper, Ray et al. (2015) suggest a representation of attention in which frontal and parietal cortices induce top-down effects on the visual, auditory and somatosensory cortices, in accordance with the majority of the studies reviewed ^65^. This frontal-parieto-occipal network encompassed the superior parietal lobe, frontal eye fields and middle frontal gyrus ^66,67^. While the dynamics of such an attentional frontal-parietal network are still debated, some studies have evaluated the cortical networks involved in the control of auditory attention using EEG source imaging methods. These studies showed a frontal lobe activation preceded by activities in multiple parietal lobe regions ^68,69^. Altogether, this is in accordance with our model suggesting a parieto-occipital activation preceding the frontal top-down mechanism.

**Figure 5.**
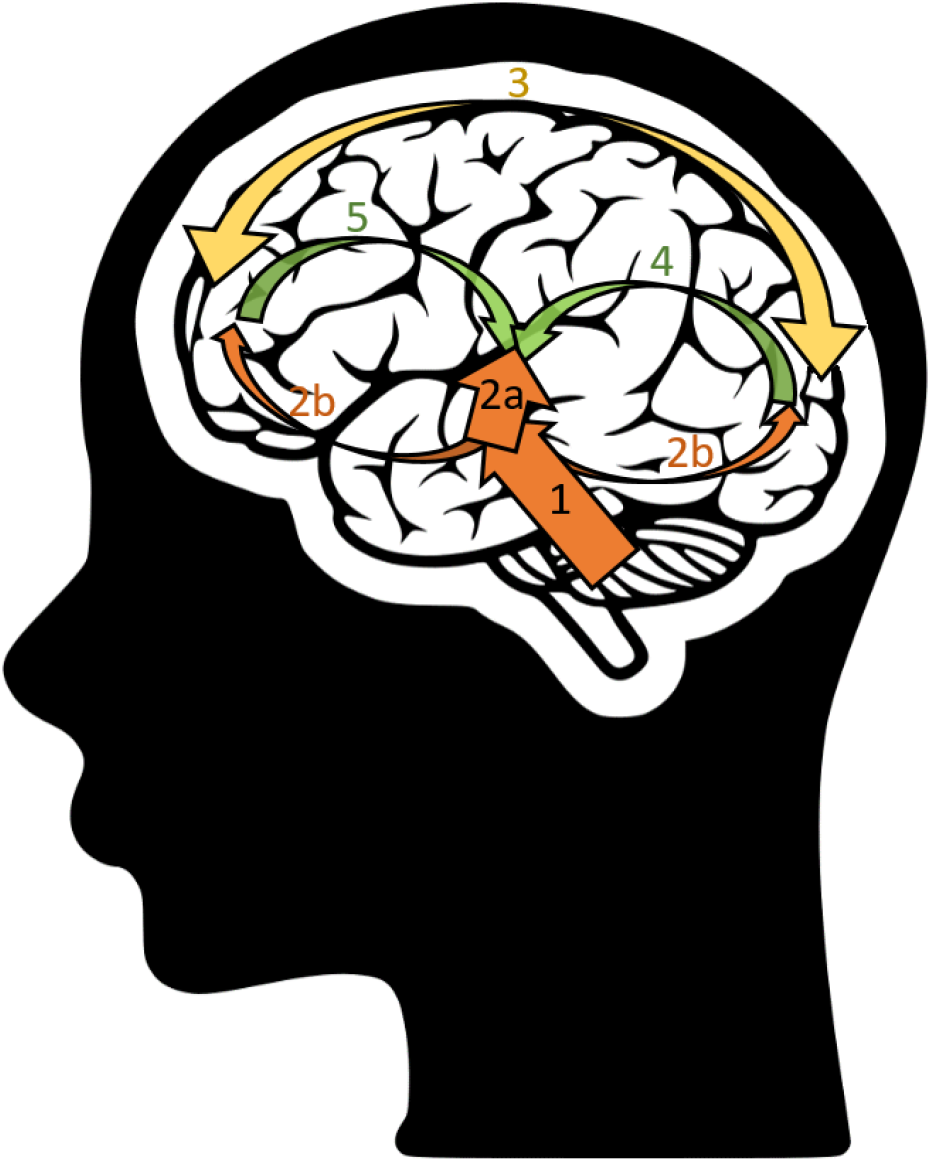
Bottom-up (in orange) and top-down (in green) processes involved in the cortical tracking of the speech envelope. First, the speech stimulus enters the ears and its information is processed all the way through the first stage of the auditory cortex (arrow 1). Once this information reaches the cortical layers, the cortical speech tracking starts. Then two streams are initiated: 2a. a bottom-up *auditory* stream (see the 0-100ms of the temporal model in Figure 4); 2b. a bottom-up *attentional* stream, activating the occipital and the frontal areas. The parieto-occipital and frontal areas interact in the dorsal and ventral attention network (arrow 3). Finally, the frontal and parieto-occipital areas impact the later-latency auditory processes through top-down processing (see latencies between 150 and 300 ms in Figure 4): the parieto-occipital (arrow 4; see the red circles in Fig.1 in the Appendix) top-down control here preceeds the frontal (arrow 5 see the green circles in Fig.1 in the Appendix) top-down control.

We do not have a clear explanation why an increase in attention induced a delay in the brain responses (see peaks in Figure 4). It has been shown that peaks from the temporal response function could be delayed when decreasing the level of speech understanding ^11,70^. Moreover, a delayed brain response to auditory stimuli has been observed in patients with decreased level of consciousness as compared to healthy volunteers ^71,72^. We would expect that a decrease in attention would produce similar results as a decrease in the level of consciousness (i.e., both would lead to a delayed auditory response), which was not the case (i.e., the peaks in the model of the “high attention” condition are delayed as compared to the peaks in the “low attention” condition). Therefore, as consciousness and sustained attention inversely affected delays in auditory response, we could argue that different processes are involved.

Our method of measuring spectral entropy and using the results to control for attention has a number of important applications. Implementing the method in real-time could allow the experimenter to act directly on the participant with the benefit of shorter experimental time with an equivalent data quality. Indeed, we here show that 20-minutes of data with high attentional level provides better performance than the standard “use-it-all” 40-minutes training dataset. This spectral entropy-based index of the participant’s attention could be used in many active attentionally-driven paradigms and correct the recorded data so the modulation of attention does not directly impact the extracted cortical measures. This is particularly relevant in neuroscience experiments in which data collection is getting longer and longer, leading to increased participant’s tiredness and attention fluctuation over time. Our results highlight the importance of controlling for attention: a large effect was found, even though the task (listening to a story) was engaging and pleasant, the participants received clear instructions to sustain attention, and compliance was monitored by asking questions.

## Supporting information

Supplementary results

## Acknowledgement

The authors thank all participants for their patience and willingness to join this study, and the authors are grateful to Eline Verschueren for her help in recruiting participants and obtaining the data. Financial support was provided by the KU Leuven Special Research Fund under grant OT/14/119 to Tom Francart. This project has received funding from the European Research Council (ERC) under the European Union’s Horizon 2020 research and innovation program (grant agreement No 637424, ERC starting Grant to Tom Francart).

## Author contributions

DL analyzed and interpreted the data, wrote the manuscript. TF designed the protocol, contributed to the writing and revised the manuscript.

## Availability of materials and data

Due to ethical concerns, supporting data cannot be made openly available. Further information about the data and conditions for access are available via Professor Tom Francart (see corresponding author).

