## Supplementary results for "The interplay of top-down focal attention and the cortical tracking of speech"

**Lesenfants D[1], Francart T[1]**

[1]*Experimental Oto-Rhino-Laryngology, Department of Neurosciences, KU Leuven, Belgium.*

#### APPENDIX

### WHICH ELECTRODES ENCOMPASS ATTENTIONAL INFORMATION?

#### Introduction

In our study, we have two neural processes working in parallel: the cortical speech tracking, computed as the correlation between the reconstructed and actual speech envelope, and the level of sustained attention, computed using spectral entropy measures over the scalp. The main analysis aims to show how the second measure (i.e., the level of sustained attention) influences the first measure (i.e., the cortical speech tracking). Those results are shown in Fig. 2-5 in our main manuscript.

In this Appendix, we aim to confirm the channel selection for the calculation of the level of sustained attention : we need to summarize the sixty-four spectral entropy time courses into a single time course representing the attentional level. In the literature frontal and parieto-occipital electrodes are typically used (Coull, Frith, Frackowiak and Grasby, 1996; Corbetta and Shulman, 2002; Moore and Armstrong, 2003; Fedorenko, Duncan and Kanwisher, 2013; Bichot, Heard, DeGennaro and Desimone, 2015).

#### Methods

To highlight electrodes involved in the attentional process, we propose a data-driven method to select electrodes by applying a linear model using ridge regression with a time-lag window of -150 to 650 ms between the cortical speech tracking and the spectral entropy values extracted at each channel location. Suppose a cortical speech tracking over time c(t) and the spectral entropy measure at each channel location SpE(t,n). We then find a linear mapping "$g_{SpE}$" allowing to understand how the changes in SpE at the different electrodes induce changes in c(t). This could be expressed by:

$$c(t) = \sum_{n} \sum_{\tau} SpE(t+\tau).\, g_{SpE}(\tau, n)$$

with an integration of spectral entropy values over a range of time lags $\tau$ (here, from -150 to 650 ms).

For each cross-validation (n = 11 folds), a decoder was first trained using 40-min of data, similarly to the main investigation. In the current investigation, the 4-min testing set was split into consecutive overlapping 1s-epochs (i.e., epochs overlap from all but one – the slide increment - sample; hop size of one sample) and a measure of cortical speech tracking c(t) was computed on each epoch as the spearman correlation between the reconstructed and actual 1s-envelopes. This results in a time series of values representing the cortical speech tracking over the testing dataset. We computed the time series for the different 4-min testing datasets (i.e., there are 11 folds, so this results in 11 time series, each time series containing a correlation value per 1s-epoch of the 4-min testing set) and then concatenated them. In parallel, we computed the entropy values at each channel location for each 1s epoch, SpE(t). We finally derived a cortical speech tracking-attention mapping, "$g_{SpE}$", by linearly combining the spectral entropy values at each channel location and their time shifted versions in order to optimally reconstruct the cortical tracking time series.

#### Results

When investigating the relation between spectral entropy and cortical speech tracking, a parieto-occipital activation (see Figure Ap1, red circles) could be observed at around 150 ms, followed by a frontal activation at around 300 ms (see Figure Ap1, green circles). This suggests that the fluctuation of the spectral entropy over these areas induces changes in the cortical tracking of the speech envelope over time. This is in accordance with the literature suggesting a network of frontal and parietal cortical areas is involved in the selection required for top-down attention.

*#Insert Figure Ap1 here*

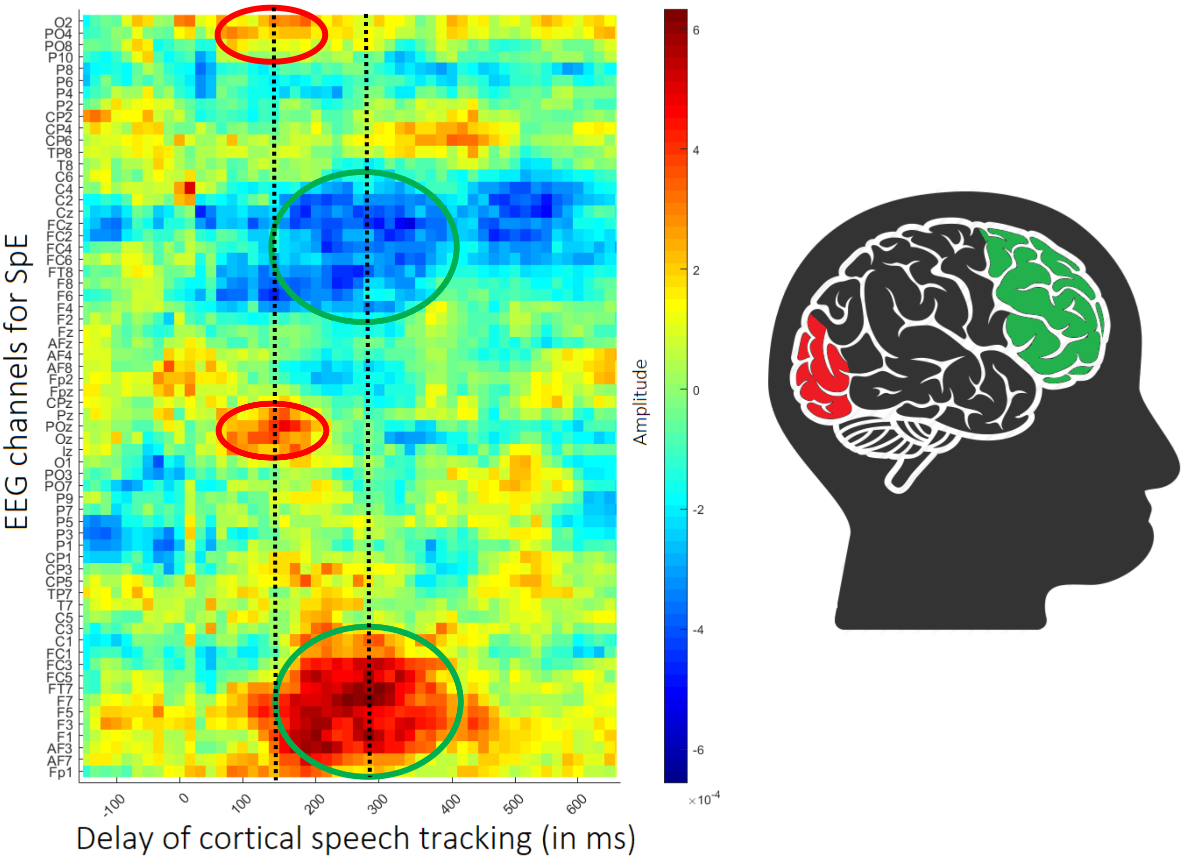


Figure Ap1 – Impact of the spectral entropy (8.5-32Hz) on the cortical tracking to the speech (0.5-8Hz). Note the activation of the occipital area at around 150 ms followed by an activation in the frontal area at around 300ms.
